# Impact of formate supplementation on body weight and plasma amino acids

**DOI:** 10.1101/2020.05.04.075796

**Authors:** Sandeep Dhayade, Matthias Pietzke, Robert Wiesheu, Jacqueline Tait-Mulder, Dimitris Athineos, David Sumpton, Seth B. Coffelt, Karen Blyth, Alexei Vazquez

## Abstract

Current nutritional recommendations are focused on energy, fat, carbohydrate, protein and vitamins. Less attention has been paid to the nutritional demand of one-carbon units for nucleotide and methionine synthesis. Here we investigate the impact of sodium formate supplementation as a nutritional intervention to increase the dietary intake of one-carbon units. A cohort of six female and six male mice received 125 mM sodium formate in the drinking water for three months. A control group of another six female and six male mice was also followed up for the same period of time. Tail vein blood samples were collected once a month and profiled with a haematology Analyser. At the end of the study blood and tissues were collected for metabolomics analysis and immune cell sorting. Formate supplementation has no significant physiological effect on male mice. Formate supplementation has no significant effect on the immune cell counts during the intervention or at the end of the study in either gender. In female mice however, the body weight and spleen wet-weight were significantly increased by formate supplementation, while the blood plasma levels of amino acids were decreased. Formate supplementation also increased the frequency of probiotic bacteria in the stools of female mice. We conclude that formate supplementation induces physiological changes in female mice.

## Introduction

Folic acid, vitamin B9, is an essential cofactor in one-carbon metabolism, a network of biochemical reactions mediating the synthesis of methionine and nucleotides ^(1)^. Our cells do not make folic acid and therefore we satisfy our daily demand for folic acid through our diet. The nutritional requirements for folic acid have been extensively addressed in the scientific literature ^(2)^. There is also an extensive literature covering the interaction between folic acid and vitamin B_12_. Methionine synthase requires vitamin B_12_ as a co-factor, to transfer one-carbon units from the folate pool to the methyl pool ^(1)^. Deficiencies in either folic acid or vitamin B_12_ lead to a spectrum of similar pathologies ^(3)^.

New studies highlight the need to investigate the nutritional requirements of the one-carbon units per se ^(4; 5)^. One-carbon units can be obtained from the amino acids serine, glycine and methionine; from methyl containing molecules such as choline, methanol, methylamine and creatine; and from the quintessential one-carbon precursor, formate ^(4; 5)^. The diversity of one-carbon nutritional sources may give the impression that there is no need to specify the nutritional requirements of one-carbon units. The view has been challenged by our observations of low serum formate levels in cancer patients and obese individuals compared to healthy controls ^(6)^.

Our daily demand of one-carbon units can be estimated from the uric acid content in urine. Uric acid is the product of purine catabolism and it contains two one-carbon units. Based on the daily urine excretion of uric acid we need about 1 gram of formate per day ^(7)^. This demand may be satisfied from the consumption of formate precursors contained in animal meats, dietary fibre and coffee ^(5; 7)^.

In the present study we investigate the impact of dietary formate intake on mammalian physiology. To this end we conducted sodium formate supplementation experiments in female and male mice. Our data suggests that sodium formate supplementation has an anabolic effect in female mice, with a significant increase in body weight and a significant reduction of plasma amino acid levels. Sodium formate also increased the frequency of pro-biotic bacteria in the gut of female mice. In the male cohort there is a striking variability from mouse to mouse in their response to sodium formate supplementation. We also profiled the immune system of female and male mice and noted no significant effect of sodium formate supplementation.

## Results

### Plasma formate

Sodium formate supplementation had no significant effect on the plasma formate concentration of female mice (**Fig. 1A**). Male mice drinking water alone had significantly lower plasma formate than aged matched female mice and the plasma formate concentration was increased by sodium formate supplementation (**Fig. 1A**). The latter was not significant due to the high variability across the male mice supplemented with sodium formate.

**Figure 1.**
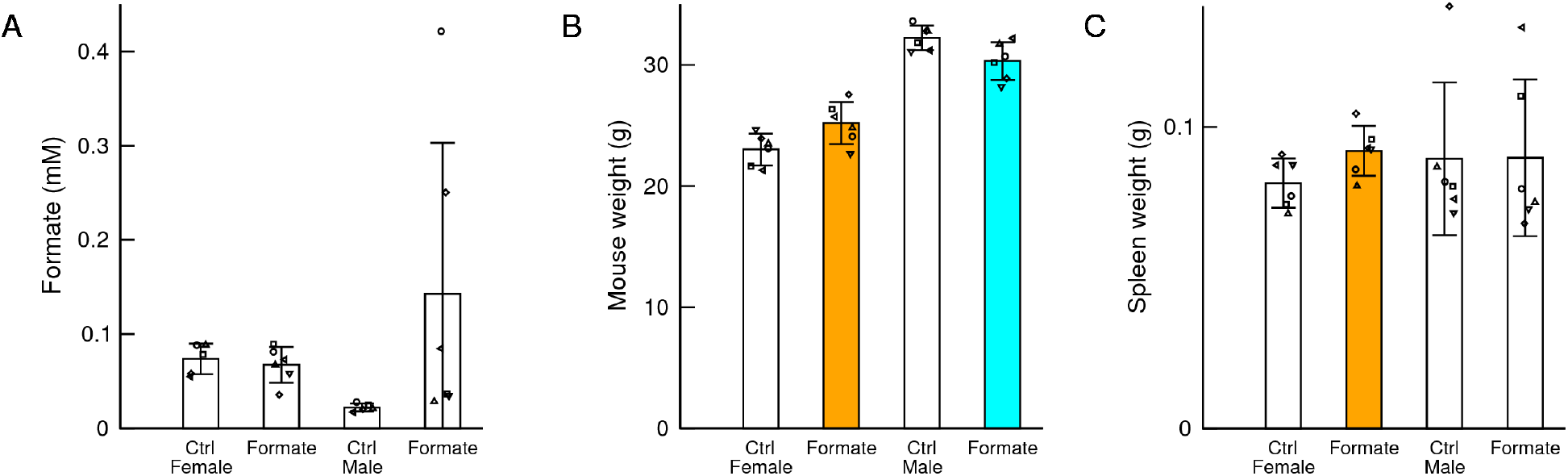
Major characteristics of the female and male cohorts drinking regular water (Ctrl) or water supplemented with 125 mM sodium formate. A) Plasma formate at the end of the study. B) Body weight at the end of the study. C) Spleen wet weight at the end of the study. Statistics: Each symbol represents a mouse, the bars heights represent the mean and the error bars the standard deviation. Coloured bars highlight a significant change (p<0.05) in the cohort with sodium formate supplementation relative to control of the same gender. Orange indicates a significant increase and cyan a significant decrease.

### Body weight

Sodium formate supplementation resulted in a significant increase in the body weight of female mice when compared to female mice drinking water alone (**Fig. 1B**). Because of the follow up immunological characterization, we also measured the wet weight of spleens at the end of the study. Sodium formate supplementation resulted in a significant increase in the wet spleen weight of female mice when compared to control female mice (**Fig. 1C**). In contrast to female mice, we observed a small but significant decrease in the body weight of male mice (**Fig. 1B**) and no significant changes in the weight of their spleens at the end of the study (**Fig. 1C**).

### Plasma metabolites

To determine which metabolic pathways might be associated with the observed body weight changes, we analysed the plasma samples using liquid chromatography and high-resolution mass spectrometry.

We performed a targeted quantification of selected metabolites (Table S1). The most striking observation was a significant reduction of all proteogenic amino acids in the plasma of female mice subject to sodium formate supplementation (**Fig. 2A**), while no significant changes were detected in the male cohorts (**Fig. 2B**). This is illustrated in **Fig. 2C,D** for the amino acids threonine and methionine. Sodium formate supplementation also increased the levels of citrate in the plasma of female mice (**Fig. 2E**). The increase in citrate suggest an increase in TCA cycle metabolites.

**Figure 2.**
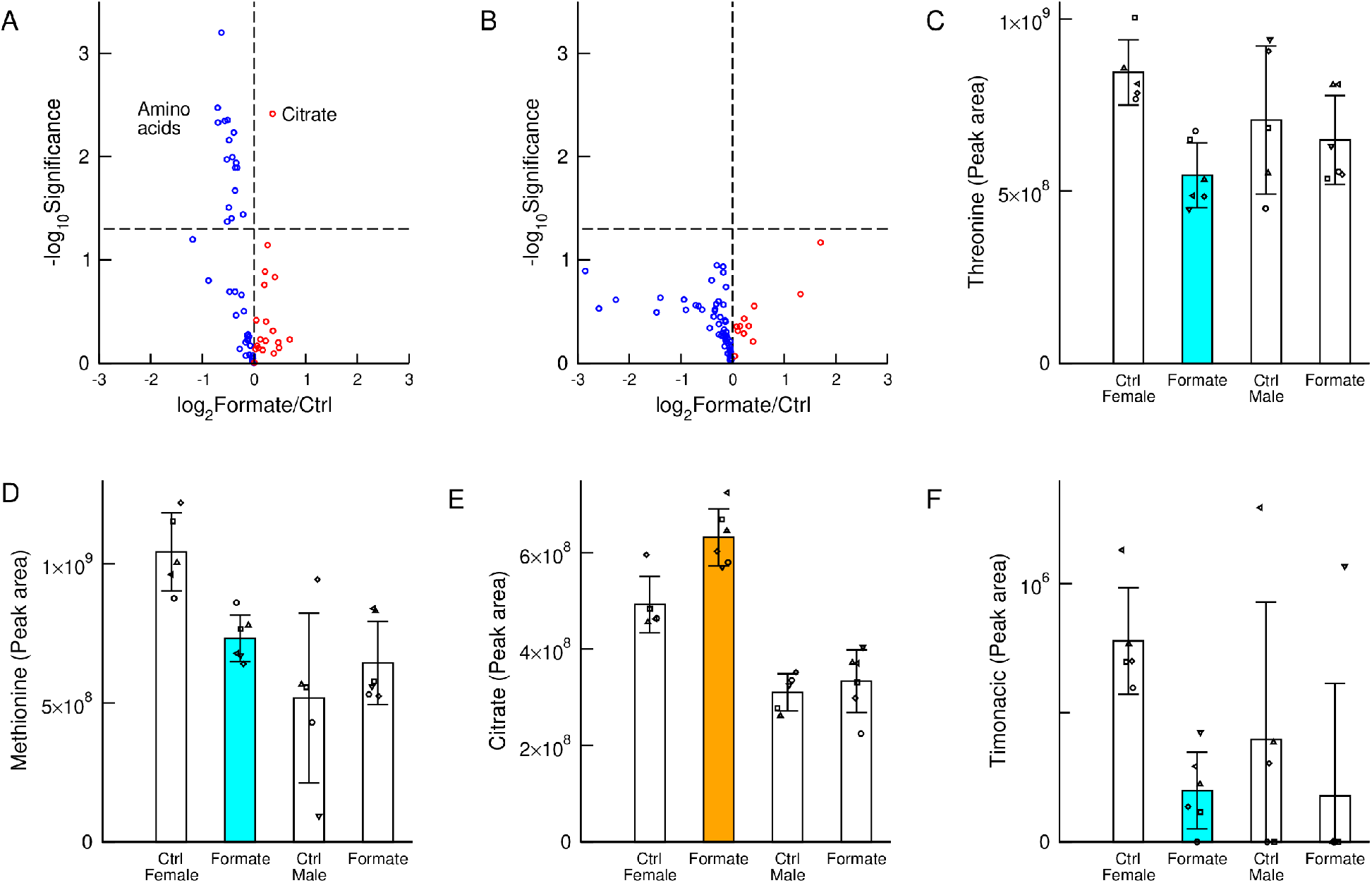
Metabolic changes. A, B) Volcano plots of metabolic differences between plasma samples of mice supplemented with 125 mM sodium formate and controls, for female A) and male mice B). The horizontal axis reports the log-ratio and the vertical axis the statistical significance (two tails and unequal variance t-test) in a logarithmic scale. C-F) Levels of the indicated metabolites across the different cohorts. Statistics: Each symbol represents a mouse, the bars heights represent the mean and the error bars the standard deviation. Coloured bars highlight a significant change (p<0.05) in the cohort with sodium formate supplementation relative to control of the same gender. Orange indicates a significant increase and cyan a significant decrease.

There was also a significant reduction of Timonacic in the plasma of female mice (**Fig. 2F**). Timonacic is an adduct formed from the reaction of formaldehyde with cysteine ^(11)^. A reduction in the plasma levels of Timonacic could be due to a reduction on the plasma levels of cysteine. Another possibility is that sodium formate induces a decrease in plasma or tissue formaldehyde and as a consequence a decrease in the levels of Timonacic, the adduct with cysteine. Since formaldehyde is a known carcinogen, these data suggest a protective effect of sodium formate supplementation with respect to cancer risk.

### Immune system

A recent study reported an age-dependent decline of the availability of one-carbon units in T cells ^(12)^. Based on this evidence we hypothesised that sodium formate supplementation may have a beneficial effect on the immune system. To obtain a longitudinal characterization of the immune system we collected tail vein blood samples once per month from each mouse and investigated whole blood leukocyte counts for both female and male cohort animals. Sodium formate supplementation had no significant effect on the counts of any immune cell subpopulation (Table S2), as shown in **Fig. 3A-D** for lymphocyte and neutrophil levels.

**Figure 3.**
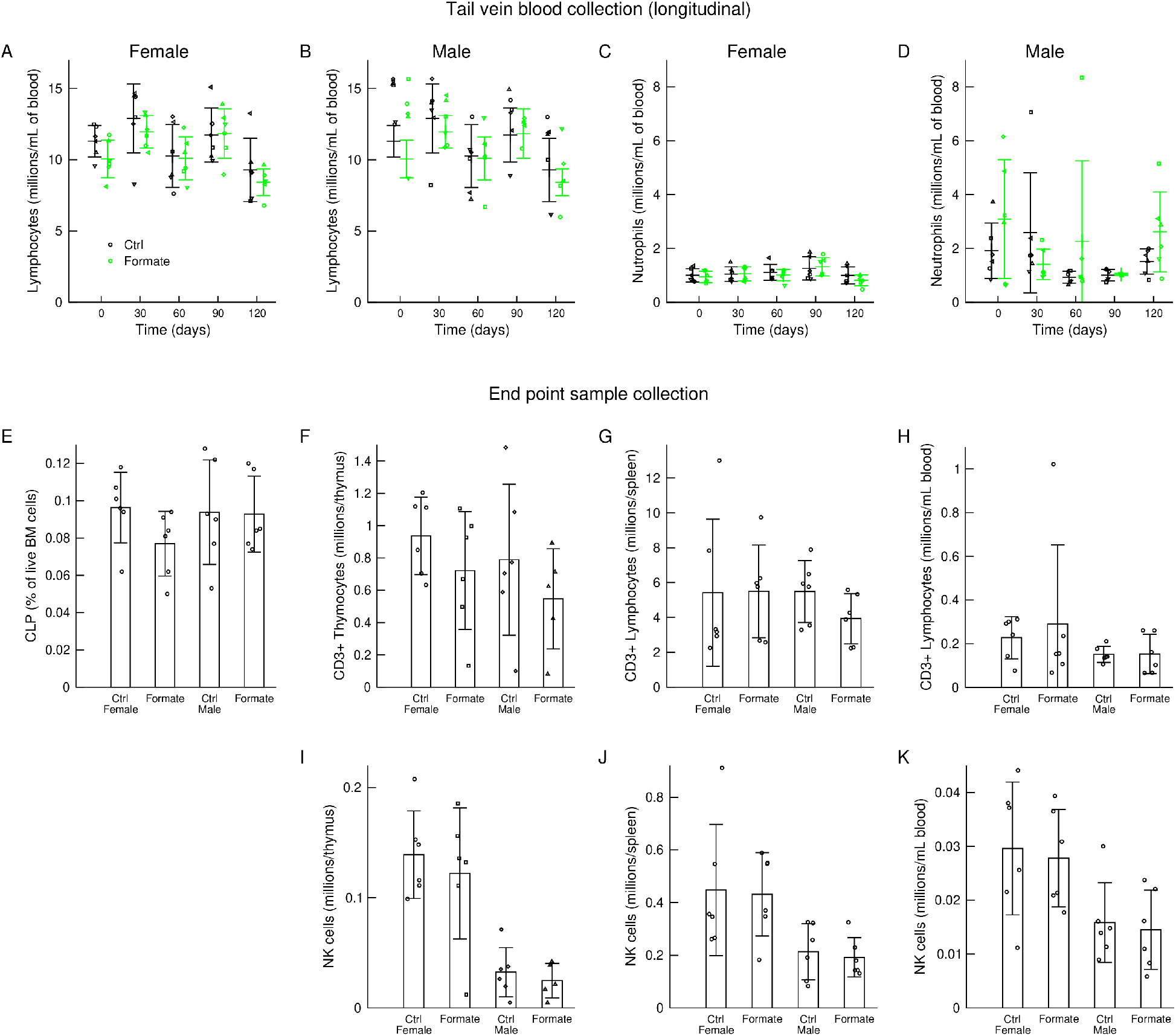
Immune system characterization. A-D) Longitudinal counts of immune system subpopulations in blood samples from the tail vein. E-K) Immune cell counts from samples and at the end of the study. Statistics: Each symbol represents a mouse, the bars heights represent the mean and the error bars the standard deviation.

We performed a more extensive characterisation of the immune system at endpoint, including blood samples and tissues from the bone morrow (BM), the thymus and the spleen. Except for a decrease of common lymphoid progenitor cells (CLPs) in the bone marrow of female mice supplemented with sodium formate (**Fig. 3E**), sodium formate supplementation had no other significant effect on the amount of major lymphocyte populations (CD3^+^ and NK cells) in the bone marrow, thymocytes, spleen lymphocytes or blood lymphocytes (**Fig. 3E-K**). We furthermore investigated the absolute numbers of distinct T cell subpopulations, including CD4^+^, CD8^+^ and γδT cells as well as the numbers of NK cell subsets based on their maturity. This analysis revealed similar numbers of each cell subset between the control group and formate-treated mice (data not shown). Of note, NK cell numbers were significantly higher in female mice compared to male mice in the thymus and spleen, but these differences remained unaffected by formate supplementation (**Fig. 3I-K**). In addition to absolute number quantification, we scrutinised the activation status of T cell and NK cell subsets, based on the expression of CD44 and CD69 alongside their cytokine production (IL-17A, IFNγ and Granzyme B). We detected the same levels of activation marker expression and cytokines for all populations between control groups and sodium formate-supplemented groups (data not shown).

### Microbiome

Formate is a central molecule in the metabolism of the gut microbiome. Formate is a by-product of the anaerobic fermentation by gut bacteria ^(13)^ and it can be also a substrate for oxidative phosphorylation by pathogenic bacteria like *E. coli* ^(14)^. Based on this evidence we conducted a characterisation of the gut microbiome.

Stool samples were collected at end point and sent to CosmoID for microbiome profiling. Sodium formate supplementation caused significant changes in the phylogenetic tree of the females microbiome but not of the male’s microbiome (**Fig. 4A,B**). At the family level, sodium formate supplementation increases the frequency of *Bifidobacteriaceae* in the stools of female mice but not of male mice (**Fig. 4C**). Since bifidobacteria are probiotic bacteria with a positive impact on health, these data indicats that sodium formate supplementation has a positive effect in the microbiome of female mice.

**Figure 4.**
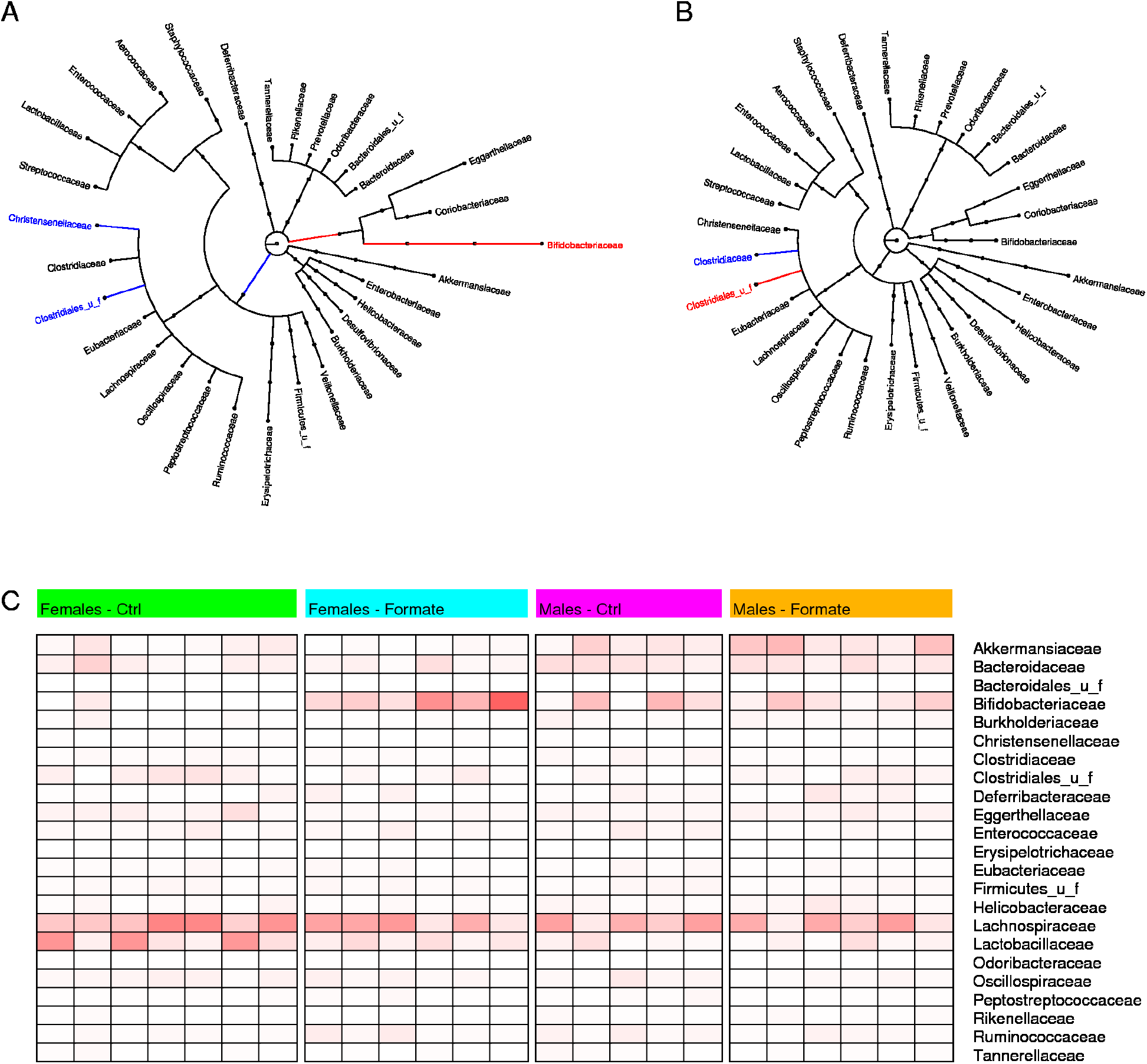
A,B) Phylogenetic tree of the identified microbiota truncated at the family level, from the stools of female (A) and male (B) mice. The longer the edges the more significant is the difference of the microbe frequency between the cohort supplemented with 125 mM sodium formate and controls. The Red colour highlights a significant frequency increase by sodium formate supplementation and the blue colour a significant decrease. C) Heatmap reporting the microbiota frequencies at the family level.

## Discussion

We have presented evidence that sodium formate supplementation induces physiological changes in a gender specific manner. In the case of females, sodium formate supplementation induces an increase in body weight, a concomitant reduction of plasma amino acids and an increase of the frequency of bifidobacteria in the gut microbiome. In contrast, we did not detect any significant effect of sodium formate supplementation in male mice.

We do not know what are the gender specific factors modulating the impact of sodium formate supplementation. One possibility is that female mice have a higher one-carbon units demand than male mice and that the amount of one-carbon units they receive through the standard chow diet is below the demand. In mammals circulating formate is at its highest levels during pregnancy ^(15; 16)^. The female mice in the cohorts investigated here were not pregnant, but they were in their reproductive age.

One aim of this study was to investigate the impact of sodium formate supplementation on the immune system. We were surprised to see no significant effect. Lymphocytes isolated from aged mice exhibit an impaired activation due to a decline in their capacity to catabolize serine to formate *in vitro*, which can be rescue by formate supplementation ^(12)^. We should bear in mind that our analysis is restricted to the immune system of antigen-naïve mice. We cannot exclude that sodium formate supplementation may have an impact on the immune response in the context of infection or nutritional deficiency.

The extrapolation of these observations to humans requires some discussion. There is evidence of ocular toxicity and death associated with high levels of formate in the context of methanol intoxication ^(17; 18; 19)^. In contrast, a few grams of formate per day are safe ^(7)^. A study supplementing healthy women with 1,300 milligrams of calcium formate three times a day for 14 days reported no evidence for ocular toxicity ^(20)^. We set the supplementation to 10% the daily one-carbon demand of mice. A similar calculation can be done in humans based on the concentration of uric acid in the urine, obtaining a daily demand of 1 gram/day or 20 mmol/day ^(7)^. The formate supplementation dose of 10 % the human daily demand translates to 2 mmol formate per day, or 136 milligrams of sodium formate per day. This dose is safe based on the toxicity study cited above. Therefore, there is no objection for the recapitulation of this study in humans.

## Experimental methods

### Ethical statement

*In vivo* experiments were carried out in dedicated barriered facilities proactive in environmental enrichment under the EU Directive 2010 and Animal (Scientific Procedures) Act (HO licence number: 70/8468) with ethical review approval (University of Glasgow). Animals were cared for by trained and licensed individuals and humanely sacrificed using Schedule 1 methods.

### Study design

Twelve female and twelve male mice where followed up for approximately 4 months, from age 56 days to age 183 days. Six female and six male mice received water with 125 mM sodium formate (Sigma) and the remaining six female and male mice drank water alone. Mice were randomized in individual groups (different gender and treatment arms).

### Experimental procedures

We first estimated the daily demand of formate for mice. We assumed the urinary excretion of allantoin as the major sink of one-carbon units in mice. Under this assumption the reference daily intake of formate for mice is *r*=2*C*_*A*_*U*/*M*_*A*_, where *C_A_* is the concentration of allantoin in mouse urine, *U* is the typical mouse urine volume per day and *M*_*A*_ is the molar mass of allantoin. The concentration of allantoin in the mouse urine is about 3.5 mg/ml ^(8)^. We assumed a urine output of 1 ml/day. The molecular mass of allantoin is 158 g/mol. Based on these numbers we calculated a demand of 40 mmol formate/day.

If we distribute the estimated 40 mmol formate in the typical daily intake of water for our C57BL/6 mice (30 millilitre), we obtain a target concentration of 1 M. To err on the safe side, we carried out a pilot study supplementing sodium formate at concentrations of 125, 250 and 500 mM in the drinking water. Mice receiving the 500 mM dose lost weight and therefore we stopped that arm of the pilot study. Mice receiving the 125 and 250 mM doses did not manifest any appreciable change in body weight. Based on this evidence we decided to use the 125 mM dose of sodium formate in our study, which represents about 10% of the daily formate demand for mice.

### Experimental animals

We use mice as a model to investigate the impact of long term formate supplementation on mammalian physiology. This choice was motivated by a number of factors. First, and most important, mice formate metabolism is representative of mammalian formate metabolism and has been proven to provide insights into the formate metabolism of mammals, including humans. Second, the C57BL/6 mice used in this study have a homogeneous genetic background and they were kept under the same conditions and diet. With this choice we thus avoid several con-founding factors that could be acting if the study is conducted in the human population. Finally, we aimed to investigate changes in the immune system at different levels, from the thymus and spleen compartments to the blood. This investigation is unfeasible in humans.

Wild-type female and male C57BL/6J mice (8 weeks of age) were purchased from Charles River.

### Housing and husbandry

Animals are housed in a purposely built facility proactive in environmental enrichment in individually ventilated cages (Techniplast) housing 3-5 animals. Animals are cared for by highly skilled technicians. For this study animals were purchased from a recognised breeder Charles River, UK. We use a 12 light/dark cycle (7am-7pm). All animals undergo a daily welfare assessment in accordance of the Standard Conditions of the Establishment Licence issued by the UK government. In addition, the researcher carried out experimental health checks (2-3 times weekly during the study).

### Sample size

In a pilot experiment investigating the effect of formate on 5 mice and 3 controls, we observed blood lymphocyte counts (millions/mL) of 6 for control mice and 10 for mice supplemented with formate, with a standard deviation of 2. Based on these numbers, a statistical power of 80% and two-sided significant level of 0.05, the estimated sample size is 4 for each group (calculated with the online tool at https://www.stat.ubc.ca/~rollin/stats/ssize/n2.html). Here we used 6 mice per group.

### Allocating animals to experimental groups

Mice were randomized in individual groups (different gender and treatment arms). All mice were of the same age.

### Experimental outcomes

Tail vein blood samples were taken once every month. Samples were analysed on an IDEXX ProCyte Dx Haematology Analyser.

At experimental endpoint, mice were sacrificed humanely by asphyxation. Blood was immediately taken by cardiac puncture, transferred into Eppendorf tubes and centrifuged at 4 °C for 10 minutes at 13k G. The supernatant was transferred into new Eppendorf tubes and flash frozen in liquid nitrogen. The spleen was weighed. Bone marrow was harvested by flushing one femur per specimen with ice-cold PBS and kept on ice until further procession. Tissues were split in half and then harvested in Eppendorf tubes and flash frozen in liquid nitrogen for metabolomics. For further analysis by multiparameter flow cytometry, samples were kept in PBS on ice. Flash frozen tissue samples were blinded with random IDs and processed by a different person for metabolite extraction and analysis. After final data analysis IDs were uncovered.

Plasma formate was measured as a benzyl-derivative. An Agilent 7890B GC was used for the measurements, equipped with a Phenomenex ZB-1701 column (30 m × 0.25 mm × 0.25 μm) coupled to an Agilent 7000 triple-quad-MS. The temperature of the inlet was 280°C, the interface temperature was 230°C, and the quadrupole temperature was 200°C and the EI voltage was set to 60 eV. 2 μl of the sample was injected and transferred to the column in split mode (1:25) with a constant gas flow through the column of 1 ml/min. The oven temperature started at 60 °C, was held for 0.5 min, followed by a ramp of 38 °C/min to 230 °C, which was held for another minute. Total run time was 6 minutes; the retention time of benzyl-formate was 3.8 min. The mass spectrometer was operated in selected ion monitoring (SIM) mode between 3.0 and 4.3 min with SIM masses of 136, 137, and 138 for M0, M+1, and M+2 (internal standard) formate, respectively. Recorded data were processed using the MassHunter Software (Agilent). Integrated peak areas were extracted and used for further quantifications using in-house scripts, including the subtraction of natural isotope abundances, as described in ^(9)^ for formate or in the supplementary materials for formaldehyde.

Plasma metabolite extraction for LC-MS analysis was performed as previously described ^(10)^. LC-MS analysis was performed as described previously using pHILIC chromatography and a Q-Exactive mass spectrometer (Thermo Fisher Scientific). Raw data analysis was performed using TraceFinder (Thermo Fisher Scientific) software.

For the Flow-cytometry immune cell analysis, single-cell suspensions of all tissues were prepared by mashing samples through 70μm cell strainer. Erythrocytes were lysed with ammonium chloride lysis buffer (10X RBC Lysis Buffer, eBioscience). For T cell activation and cytokine staining, cells were stimulated with Cell Activation Cocktail with Brefeldin A (Biolegend, 1:500) in IMDM supplemented with 8% FCS, 0.5% 2-Mercaptoethanol and penicillin-streptomycin solution (Sigma Aldrich)) for 3h at 37°C. Single-cell suspensions were further incubated with TruStain FcX (Biolegend) for 20min at 4°C to block Fc receptors, followed by incubation with fluorochrome-conjugated antibodies for 30min at 4°C in the dark in brilliant stain buffer (BD Biosciences). Prior to fixation and permeabilization for intracellular staining with Cytofix/Cytoperm (BD Bioscience), cells were stained with Zombie NIR Fixable Viability kit (Biolegend) (1:400, in PBS) for 20min at 4°C in the dark for dead cell exclusion. Fluorochrome-conjugated antibodies for intracellular epitopes were diluted in permeabilization buffer and incubated for 30min at 4°C in the dark. Samples were acquired using BD LSR II flow cytometer and Diva software. Data analysis was performed by using FlowJo version 9.9.6 and fluorescence minus one (FMO) controls to facilitate gating. Gating strategies for respective immune cell populations can be seen in supplementary figures 1–2. For calculations of absolute numbers, 123count eBeads (eBioscience) were used according to manufacturer’s instructions.

#### Antibodies used for spleen, thymus and blood samples

CD19 (eBioscience, clone 1D3, APC-eFluor780, 1:400), EpCAM (eBioscience, clone G8.8, APC-eFluor780, 1:100), CD3 (Biolegend, clone 17A2, BV650, 1:100), CD4 (Biolegend, cloneRM4-5, BV605, 1:100), CD8 (BD Bioscience, clone 53-6.7, BUV395, 1:100), TCRδ (eBioscience, clone GL3, FITC, 1:200), CD11b (eBioscience, clone M1/70, BV785, 1:800), CD27 (Biolegend, clone LG.3A10, PE-Dazzle594, 1:400), CD44 (Biolegend, clone IM7, PerCP-Cy5.5, 1:100), CD69 (Biolegend, clone H1.2F3, BV510, 1:50), NK1.1 (Biolegend, clone PK136, BV421, 1:50), IFNγ (eBioscience, clone XMG1.2, PE-Cy7, 1:200), IL-17A (eBioscience, clone eBio17B7, PE, 1:100), Granzyme B (Biolegend, clone GB11, AF647, 1:50).

#### Antibodies used for bone marrow analysis

CD4 (Invitrogen, clone GK1.5, APC-eFluor780, 1:100), CD8 (Invitrogen, clone 53-6.7, APC-eFluor780, 1:100), CD11b (eBioscience, clone M1/70, APC-eFluor780, 1:800), CD19 (eBioscience, clone 1D3, APC-eFluor780, 1:400), Ter-119 (eBioscience, clone TER-119, APC-eFluor780, 1:50), CD11c (Biolegend, clone N418, PerCP/Cy5.5, 1:800), CD3ε (eBioscience, clone 145-2C11, FITC, 1:100), Ly-6A/E (Sca-1) (Biolegend, clone D7, BV605, 1:100), NK1.1 (Biolegend, clone PK136, BV421, 1:50), CD117 (c-Kit) (Biolegend, clone 2B8, APC, 1:50), CD127 (IL-7Rα) (Biolegend, clone A7R34, PE, 1:100).

For the microbiome profiling, stool samples were collected at endpoint and stored in Eppendorf tubes containing DNA/RNA shield (Zymo Research). Samples were sent in dry ice to CosmoID for 16S sequencing and inference of microbiome abundance.

### Statistical methods

Reported Statistical significances were determined using a t-test with two tails and unequal variance.

## Acknowledgements

We thank Catherine Winchester for helpful comments about the manuscript.

## Authorship

SD carried on the formate supplementation experiment, performed the longitudinal tail vein bleeds and the longitudinal blood samples analysis, under the supervision of KB. MP performed the metabolomic profiling with the assistance of JTM and DS. RW performed the immune system profiling of the endpoint samples, under the supervision of SC. DA conducted the pilot study to set the formate supplementation dose. AV conceived the project and performed the statistical analysis of the microbiome data. MP, RW, SC, KV and AV wrote the manuscript.

---

**Supplementary figure 1:**
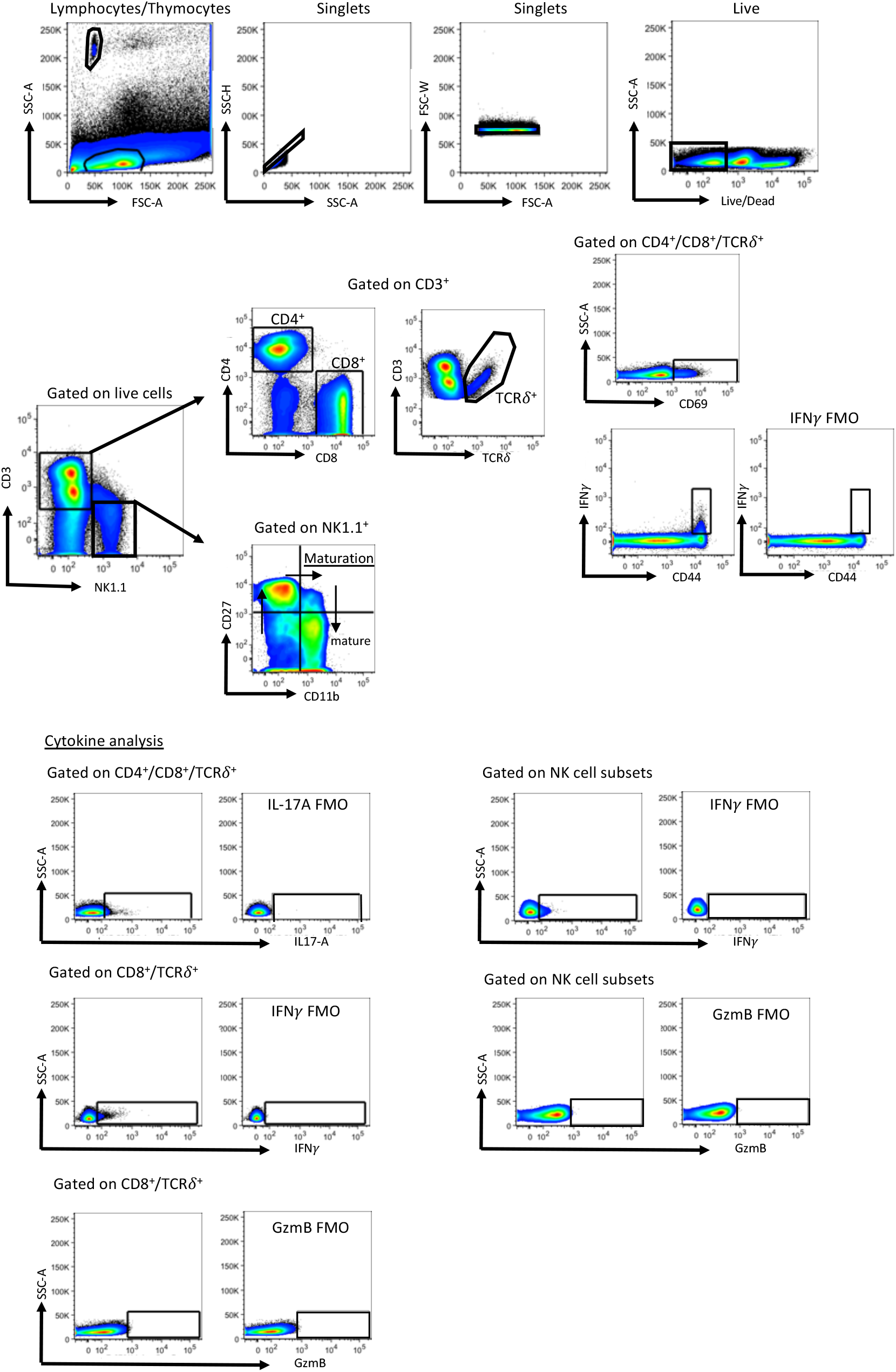
Gating strategy for various immune cell populations of blood, thymus and spleen. Lymphocytes/thymocytes were excluded for doublets twice before gating on live cells, which also excludes CD19^+^ and EpCAM^+^ cells. CD3^+^ lymphocytes and thymocytes as well as NK cells were investigated as depicted. Activation status of T cell subsets (including CD4^+^, CD8^+^ and γδ T cells) was investigated based on expression of CD69 as well as CD44 and IFNγ. NK cell maturity was assessed by CD11b and CD27 expression. NK cell subpopulations as well as T cell subsets were analysed for cytokine production as indicated. Fluorescence minus one (FMO) controls were used to facilitate gating.

**Supplementary figure 2:**
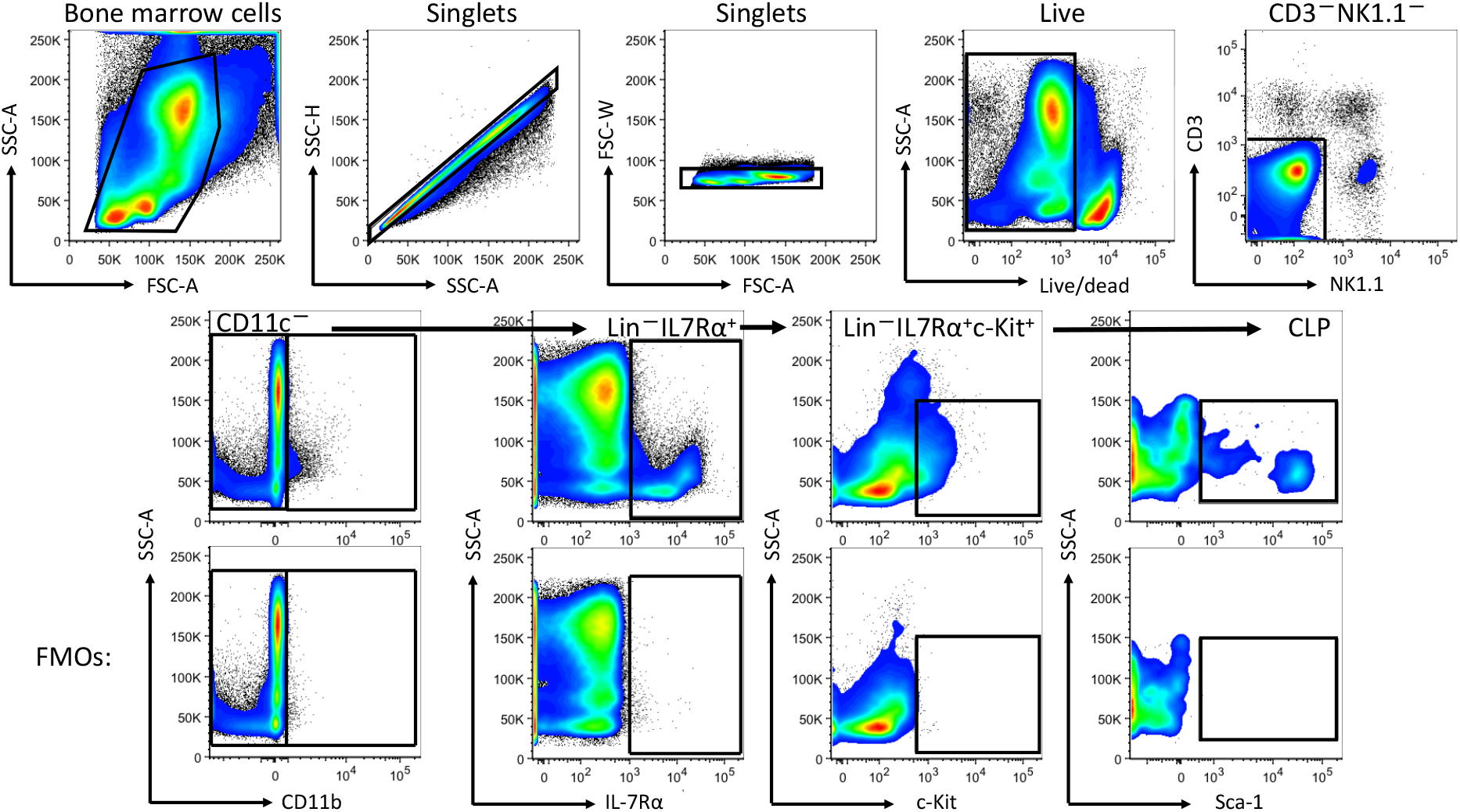
Gating strategy for bone marrow analysis. Bulk bone marrow cells were excluded for doublets twice before gating on live cells, thus further excluding lineage markers such as CD4, CD8, CD11b, CD19 as well as Ter-119. CD11c^−^ cells were considered as lineage-marker negative (Lin^−^) cell population. Common lymphoid progenitor (CLP) cells were characterized as Lin^−^IL7Rα^+^c-Kit^+^Sca-1^+^. Fluorescence minus one (FMO) controls were used to facilitate gating.

